# High-density, long-lasting, and multi-region electrophysiological recordings using polymer electrode arrays

**DOI:** 10.1101/242693

**Authors:** Jason E. Chung, Hannah R. Joo, Jiang Lan Fan, Daniel F. Liu, Alex H. Barnett, Supin Chen, Charlotte Geaghan-Breiner, Mattias P. Karlsson, Magnus Karlsson, Kye Y. Lee, Hexin Liang, Jeremy F. Magland, Angela C. Tooker, Leslie F. Greengard, Vanessa M. Tolosa, Loren M. Frank

**Affiliations:** Medical Scientist Training Program and Neuroscience Graduate Program, University of California San Francisco, CA 94158, USA.; Kavli Institute for Fundamental Neuroscience, Center for Integrative Neuroscience, and Department of Physiology, University of California San Francisco, CA 94158, USA.; Bioengineering Graduate Program, University of California San Francisco, CA 94158, USA.; Center for Computational Biology, Flatiron Institute, 162 Fifth Avenue, New York, NY 10010, USA.; Center for Micro- and Nano-Technology, Lawrence Livermore National Laboratory, Livermore, CA 94550, USA.; Neuralink Corp., San Francisco, CA 94107, USA.; SpikeGadgets IIc., San Francisco, CA 94158, USA.; Howard Hughes Medical Institute; Courant Institute, NYU, New York, NY 10012, USA.

## Abstract

The brain is a massive neuronal network, organized into anatomically distributed sub-circuits, with functionally relevant activity occurring at timescales ranging from milliseconds to months. Current methods to monitor neural activity, however, lack the necessary conjunction of anatomical spatial coverage, temporal resolution, and long-term stability to measure this distributed activity. Here we introduce a large-scale, multi-site recording platform that integrates polymer electrodes with a modular stacking headstage design supporting up to 1024 recording channels in freely behaving rats. This system can support months-long recordings from hundreds of well-isolated units across multiple brain regions. Moreover, these recordings are stable enough to track 25% of single units for over a week. This platform enables large-scale electrophysiological interrogation of the fast dynamics and long-timescale evolution of anatomically distributed circuits, and thereby provides a new tool for understanding brain activity.

## Introduction

An ideal method to observe brain dynamics would monitor many neurons, have high spatial and temporal resolution, enable access to multiple distant brain regions, and be usable in awake, freely behaving subjects. Recent work illustrates the potential power of this approach in producing scientific insight: spiking activity from 100-250 simultaneously recorded units within one region can be used to discover single-event content and dynamics (Pfeiffer and Foster, 2013, 2015), activity structure that is not possible to resolve with fewer recorded neurons. Indeed, in the spatial domain, if it were possible to record from similarly high numbers of neurons from multiple brain regions, analogous discoveries in distributed neural computation and function are likely to follow. Furthermore, in the temporal domain, if it were possible not only to record at millisecond precision, but to do so continuously over the span of hours, days, and weeks, such access could yield transformative insight into neural dynamics. Here, too, previous experimental efforts suggest this possibility: recording small numbers of neurons over the span of days has revealed surprising long-timescale firing patterns with functional implications, particularly with respect to learning (Hengen et al., 2013; Hengen et al., 2016).

Most current approaches are optimized exclusively for either the spatial or temporal domain. For example, one- and two-photon imaging can provide long-lasting, cell-type specific, and stable sampling of neuronal populations, but are limited by the temporal resolution and signal to noise ratio of the indicators (Chen et al., 2013), making it difficult to infer the precise timing of single spikes *in vivo*. Further, these methods do not permit continuous (24 hours a day, 7 days a week) recordings of brain activity. In contrast, electrophysiological approaches provide excellent temporal resolution, but technologies available in awake, freely-behaving animals are generally limited in their unit yields, spatial coverage, signal longevity, signal stability, and/or adaptability across species for continuous recording. For example, the recently developed Neuropixel probe (Jun et al., 2017) allows for recordings from 384 of 960 total sites, but recording sites must be collinear, and it remains to be established whether long-term tracking of individual neurons is possible. Conversely, long term, continuous recordings of small numbers of neurons were recently documented with a 64-channel tetrode-based system (Dhawale et al., 2017), but this approach does not provide a clear path to recordings from much larger ensembles.

Here we introduce a polymer probe-based system that overcomes the limitations of currently available technologies. Polymer devices achieve the recording contact density of silicon devices with the modularity and longevity of microwires. Polymer arrays can also provide a neural interface that is biocompatible (Jeong et al., 2015; Kim et al., 2013; Lee et al., 2017a; Luan et al., 2017) and flexible enough to counteract micromotions of the array relative to the brain (Gilletti and Muthuswamy, 2006). Until now, however, polymer arrays capable of resolving single neurons had not been developed past proof-of-concept (Kuo et al., 2013; Luan et al., 2017; Rodger et al., 2008; Seo et al., 2015; Seo et al., 2016; Tooker et al., 2014; Xie et al., 2015). Our system makes it possible to measure the activity of hundreds of single neurons across multiple, anatomically distant structures in freely-behaving animals. The system furthermore supports continuous 24/7 recording and yields high quality, large-scale single unit recordings for at least five months. In conjunction with this recording system, we adapt the MountainSort (Chung et al., 2017) spike sorting system to link clustered units across time segments, demonstrating stable recordings from 25% of individual neurons for over a week.

## Results

### Modular implantation platform

Simultaneous, large-scale single-unit recording in a distributed neural circuit requires that recording electrodes be flexibly distributed across the brain, and at high enough density to yield hundreds of putative single neurons. In the past this has necessitated a choice between a few high-density arrays with rigid geometries, or many lower-density arrays (or single channels) that can be arbitrarily and precisely distributed across the brain. Our approach, outlined in Fig. 1a, reduces the need for this tradeoff, allowing for high-resolution sampling across multiple targeted regions.

**Figure 1.**
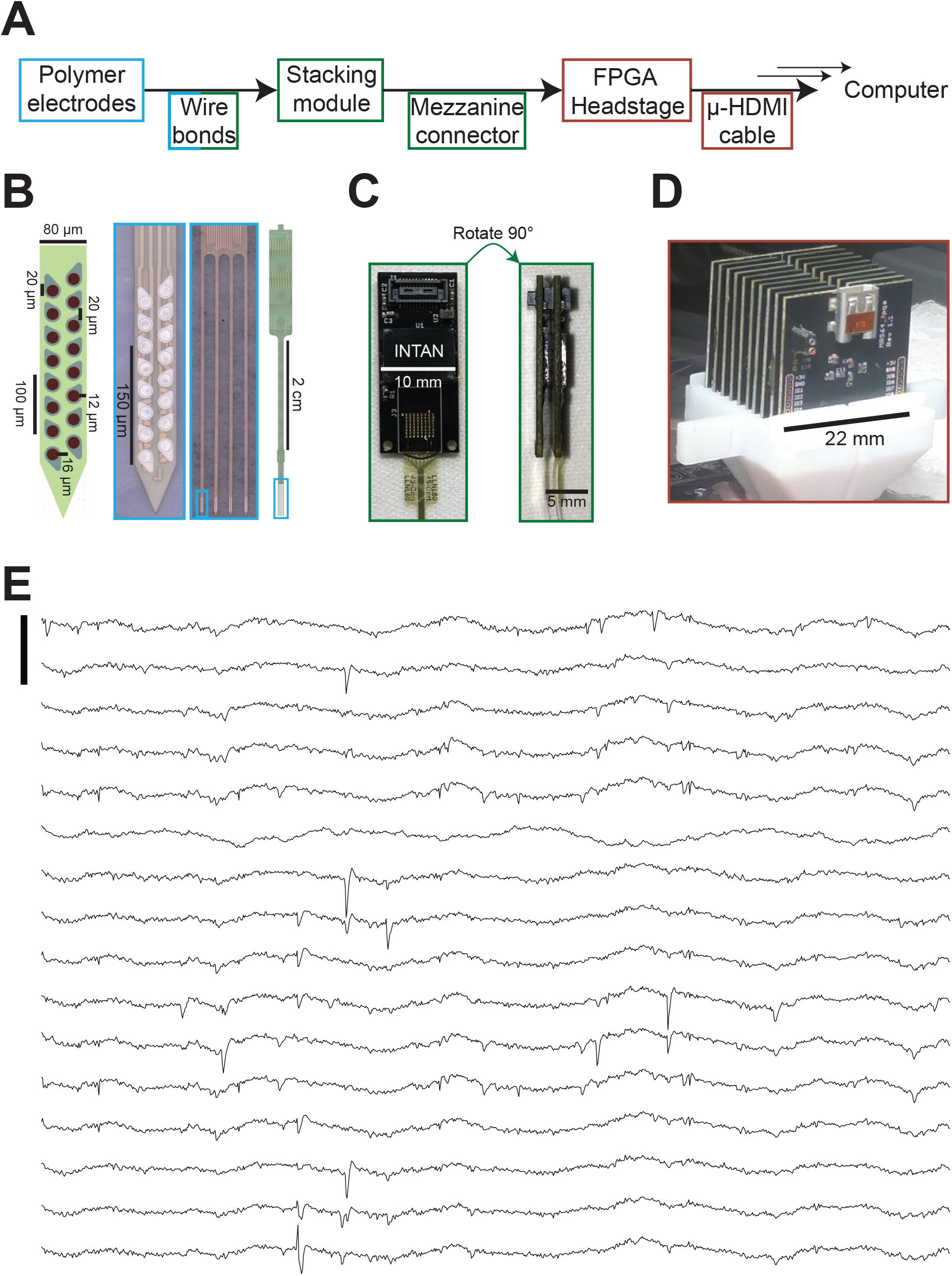
Modular 1024-channel implantation platform overview. (A) Data path from electrode to computer, with box color corresponding to related components in following subfigures. (B) Polymer electrode array. Left, schematic of 16-channel shank of polymer array designed for single-unit recording. Shank is 14 μm thick. Middle-left, image of 16-ch shank. Middle-right, 4-shank (250 μm edge-to-edge spacing), 64-channel array. Right, full polymer array, bond pads at top of array. (C) Left, view of individual 64-channel module with amplifying, digitizing, and multiplexing chip (Intan Technologies) wire-bonded onto board, and mezzanine-style connector attached at top of board. Right, two modules stacked together. (D) Full 1024-channel, 16-module, recording system stacked into FPGA headstage (SpikeGadgets llc) during implantation. (E) Raw 100 ms traces from one 16-ch shank. Scalebar corresponds to 1 mv.

Multishank polymer electrode arrays form the modular implantable unit. Each 32- or 64-channel polyimide array (Tooker et al., 2012a, b) consists of two or four shanks respectively, with 16 channels per shank. Each channel consists of a platinum electrode covered by electrically deposited PEDOT-PSS (Ludwig et al., 2006) (Fig. 1b). Each 32-channel device has an attached 32-channel omnetics connector, two of which can be accommodated by the pair of mating connectors on each printed circuit board (PCB). The PCB is wire-bonded to a 64-channel amplifying, digitizing, and multiplexing chip (INTAN technologies). Each 64-channel device is directly wire-bonded to a similar PCB. The resulting modules (Fig. 1c) can be stacked using mezzanine connectors and connected to a field programmable gate array (FPGA, SpikeGadgets LLC) which supports up to two stacks of eight modules, for a total of 1024 channels (Fig 1d). The FPGA synchronizes the modules and converts the serial peripheral interface bus (SPI) signal from each module to high-definition multimedia interface (HDMI) format. The 1024 channel, 30 KHz / channel data is streamed via a micro-HDMI cable through a low-torque HDMI commutator (SpikeGadgets LLC) and data acquisition main control unit (MCU, SpikeGadgets LLC) to the data acquisition computer where it is visualized and saved (Fig. 1e). Streaming high speed data through a commutator enables continuous recordings.

The flexibility of polyimide arrays increases biocompatibility (Lee et al., 2017a) but presents a challenge to implantation. Here we employ our previously developed insertion system, which uses a detachable silicon stiffener. Stiffener-attached arrays are inserted serially into brain tissue (Felix et al., 2013) and subsequently tethered to a custom 3d-printed base piece, which is contoured and anchored to the skull (Supplemental Fig. 1; See Methods for detailed description of the implantation procedure). Serial insertion allows multiple arrays to be placed within a single brain region (<1 mm between inserted probes). The rest of the implant is then assembled; silicone gel is added to stabilize the brain, and silicone elastomer is added to protect the polymer arrays from damage and active electronic components from moisture. The entire system is then protected with a custom 3d-printed casing and passive aluminum heatsinks for impact resistance and heat dissipation (Supplemental Fig. 1).

### Recordings of hundreds of single units distributed across multiple regions

Information processing in the brain is accomplished by the millisecond-timescale interactions of thousands of single neurons (or more) distributed across multiple regions. To demonstrate our platform’s ability to resolve network events spanning multiple regions, we examined data from an animal implanted with the full 16-module system. Of these, 8 modules were used for single-unit recording (see methods for more details). Data were collected during a rest period in a familiar environment. Spike sorting using MountainSort (Chung et al., 2017) on data from these 512 channels 45 days after implantation produced 1533 clusters with a continuum of qualities. Three-hundred and seventy-five of the 1533 clusters exceeded our previously established (Chung et al., 2017) conservative cluster quality metric thresholds (isolation > 0.96, noise overlap < 0.03), and are henceforth considered single units (Fig 2a). The modules used for single unit recording were distributed among medial prefrontal cortex (mPFC, n = 2), orbitofrontal cortex (OFC, n = 4), and ventral striatum (VS, n = 2), and polymer probes designed for recording local field potentials (LFP) were targeted to the hippocampus (HPC, n = 2) (Fig 2b).

### Coordination across multiple regions during hippocampal sharp wave-ripples

The simultaneous recording of single units across multiple regions makes it possible to examine cross-area coordination. Here we focused on times when we detected hippocampal sharp wave-ripples (SWRs). The SWR (Buzsaki, 2015) is an event of synchronous hippocampal population firing known to influence activity across the majority of the brain (Logothetis et al., 2012). These earlier studies (Khodagholy et al., 2017; Logothetis et al., 2012) leveraged methods that had large spatial coverage but were lacking in single-unit resolution. In complement, studies utilizing dual-site recordings revealed that neurons across many cortical (Chrobak and Buzsaki, 1996; Isomura et al., 2006; Jadhav et al., 2016; Ji and Wilson, 2007; Sirota et al., 2003) and subcortical regions (Dragoi et al., 1999; Lansink et al., 2009; Pennartz et al., 2004) show changes in firing rates around the time of SWRs. As a result, it remains unknown if the firing rate changes are coordinated among regions.

Changes in activity across the population of 375 single units was evident during individual SWRs (Fig. 2c, d). Across all SWRs, these changes result in significant increases and decreases in firing of a subset of units in each region (Fig. 2e). We confirmed previous reports of mPFC and NAc modulation (Lansink et al., 2009; Tang et al., 2017; Wierzynski et al., 2009): 19 of 61 mPFC (13 positively, 6 negatively) (p < 1.0e-4 as compared to expected proportion, z-test for proportions) and 27 of 118 NAc (24 positively, 3 negatively) (p < 1.0e-4, z-test for proportions) showed SWR modulation based on a p < 0.05 threshold (see methods). We also found that 28 of 196 OFC units were SWR-modulated (18 positively, 10 negatively) (p < 1.0e-3 z-test for proportions), providing a further confirmation that SWR events engage activity across many cortical regions.

**Figure 2.**
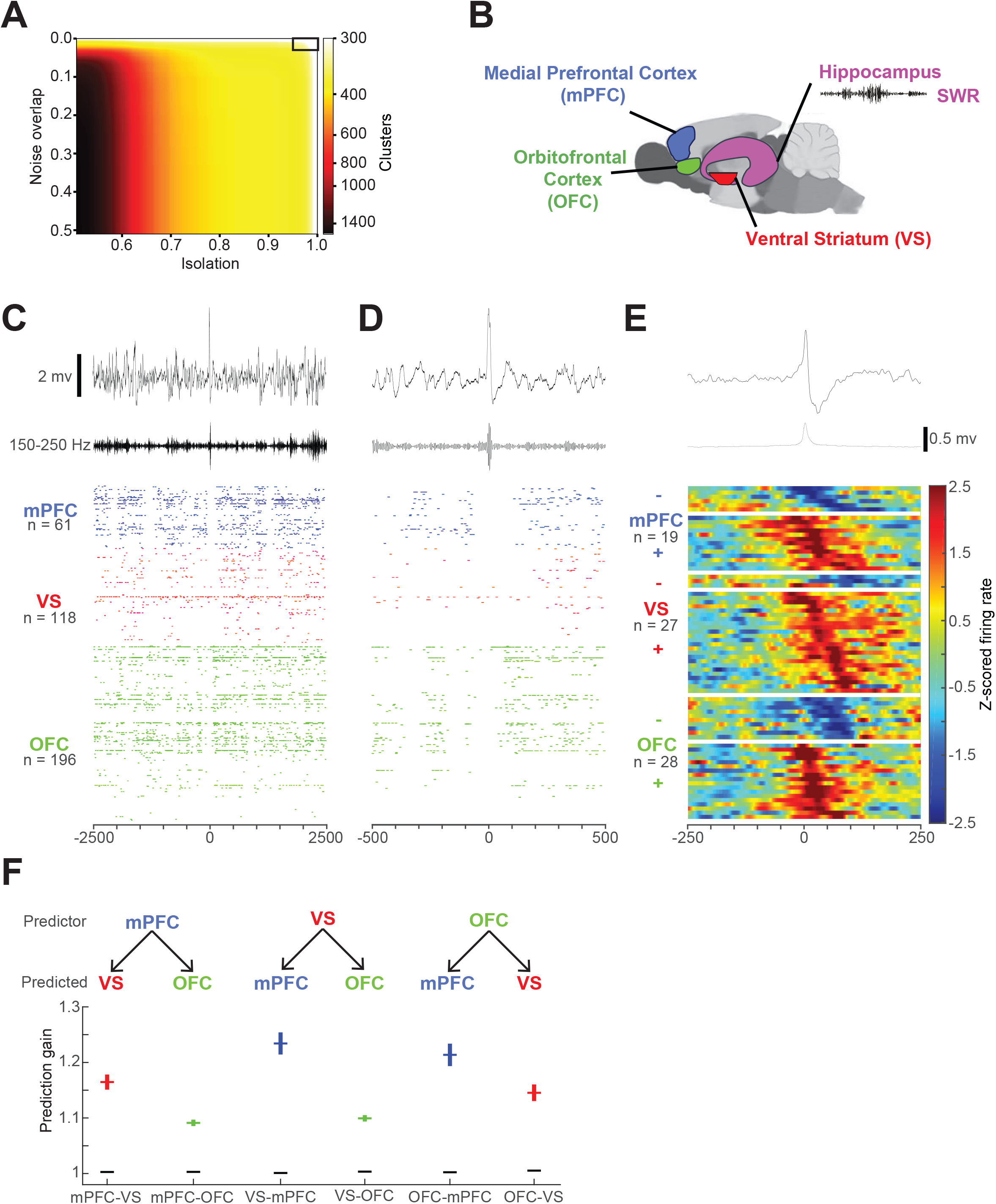
Large-scale, distributed recording. (A) Number of putative single-unit clusters from 512 channels (of the 1024-channel implant), stratified by quality metric thresholds. Automated curation using MountainSort (noise overlap 0.03, isolation 0.96, black box in upper right) resulted in the identification of 375 single units from the 512 channels. (B) Schematic of the rat brain with targeted regions highlighted. (C) Top, 5 second raw LFP trace from one of 128 channels implanted into Hippocampus, centered on a SWR. Middle, 150 – 250 Hz filtered trace. Bottom, spike rasters from 375 simultaneously recorded neurons from the same time period, with colors corresponding to the highlighted region. Horizontal axis in ms. (D) As in (C), but for 1 second centered around the same event. (E) Averaged traces for average LFP (top), power (middle, 150 – 250 Hz). Bottom, normalized firing rate, peri-SWR histograms for the significantly SWR-modulated neurons, separated by recording location, and ordered by time of trough or time of peak (calculated from 4,046 SWRs). (F) Prediction gain for each set of regions. Top, predictor region, with arrow to predicted region below. Mean prediction gain (horizontal line) ± standard error (vertical lines) for each predictor-predicted set of regions. Color of bar corresponds to each predicted region, as shown in (B). Shuffled prediction gains shown in black.

The large number of single units made it possible to show that spiking patterns are coordinated across multiple regions during SWRs. We used cross-validated generalized linear models (Rothschild et al., 2017) to determine whether ensemble firing patterns in mPFC, NAc, or OFC could significantly predict the firing rate of individual cells in the other regions at the times of SWRs (see Methods). This prediction was highly significant for all pairs of regions (prediction gains reported as mean ± standard error and p-values are from two-tailed Wilicoxon rank sum test: mPFC predicting NAc, 1.16 ± 0.01, shuffle 1.00 ± 9.8e-5, p = 1.7e-74; mPFC predicting OFC, 1.09 ± 0.01, shuffle 1.00 ± 9.1e-5, p = 8.2e-116; NAc preding mPFC, 1.23 ± 0.02, shuffle 1.00 ± 7.7e-5, p = 1.5e-38; NAc predicting OFC, 1.10 ± 0.01, shuffle 1.00 ± 1.1e-4, p = 2.1e-109; OFC predicting mPFC, 1.21 ± 0.02, shuffle 1.00 ± 3.2e-4, p = 9.8e-37; OFC predicting NAc, 1.15 ± 0.01, shuffle 1.01 ± 4.5e-4, p = 7.5e-54; Fig. 2e). Together, these findings illustrate the power of large-scale, distributed recordings and provide the first evidence of coordinated firing patterns across multiple regions during SWRs.

### Longevity of single-unit recording

While polymer devices have shown promise in achieving a long-term, biocompatible interface with neuronal tissue (Kuo et al., 2013; Luan et al., 2017; Rodger et al., 2008; Seo et al., 2015; Seo et al., 2016; Tooker et al., 2014; Xie et al., 2015), their benefits have not yet been combined in configurations and systems capable of sampling many neurons simultaneously. To evaluate the high yield single-unit recording capabilities of polymer arrays in the long term, we implanted three rats with polymer probes into mPFC or OFC for 160 days or more (one 72-ch implant, one 128-ch implant, and one 288-ch implant, see Methods).

These implants yielded long-lasting, high-quality recordings (Fig. 3a), with some initial variability across a six-week timescale, consistent with the brain’s recovery from an acute injury and the transition to a stable, chronic response (Supplementary Fig. 2). Subsequently, recording yield was stable until the end of recording (experiments terminated at 160 days to ensure the availability of histology), yielding up to 45 total units on an individual shank and ~1 single-unit per contact on average (Fig. 3a). Importantly, even after 160 days, our system continued to yield well-isolated individual single units (Fig. 3b), and in one case we extended our recordings to 283 days with only minimal decline in the number of well-isolated units (from 27 single-units at day 45 post-implant to 16 single-units at 283 days post-implant; Supplemental Fig. 2c).

**Figure 3.**
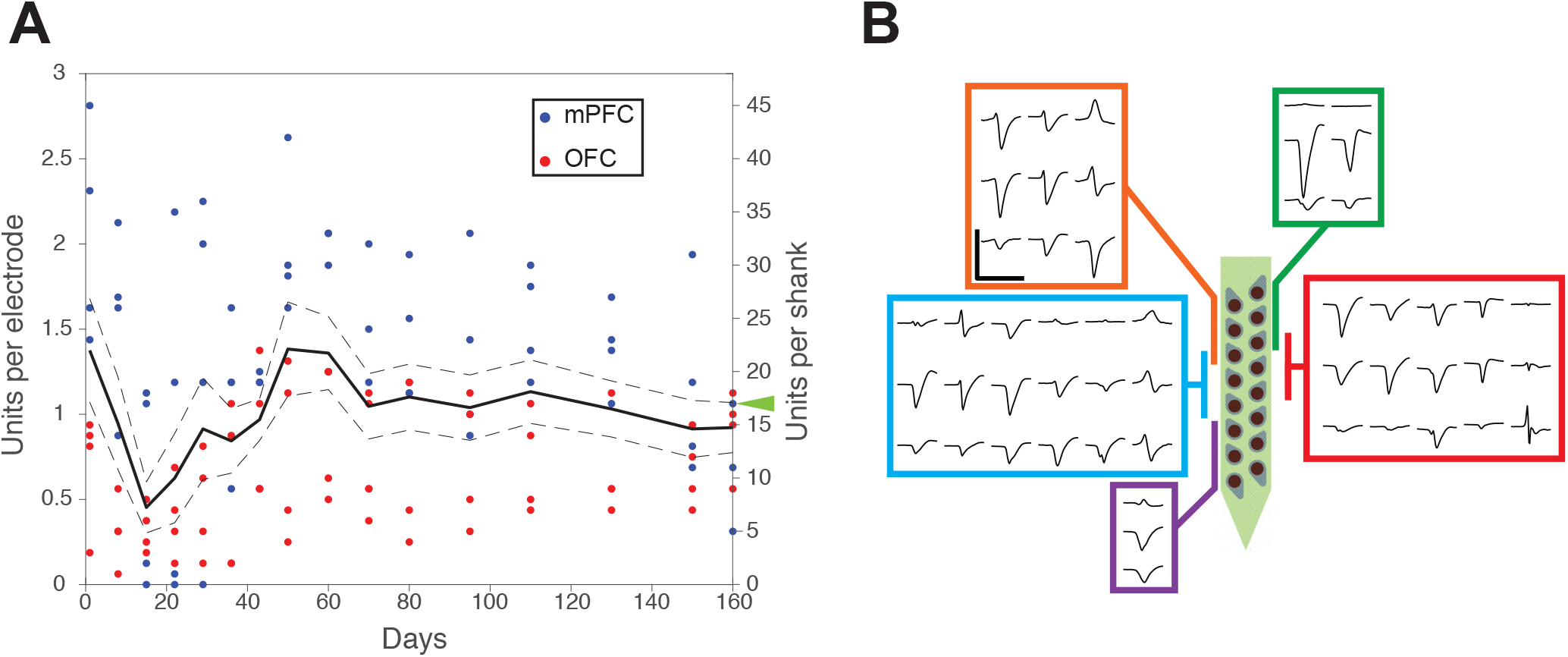
Single-unit recording yield of polymer probes over time. (A) Single-unit yields for polymer probes per channel (left y-axis) or per 16-ch shank (right y-axis) over 160 days postimplantation (x-axis) in rats. Solid line is the mean cell yield across 8 shanks, dotted lines ± 1 SE. Individual time points per shank are shown as color-coded dots by region. (B) Waveforms for units clustered for data point with green arrowhead. Scale bar corresponds to 200 μv and 2 ms.

### Stability of recording

The ability to track individual neurons across days depends upon stable recordings and a clustering strategy that is robust to changes in waveform shape resulting from electrode movement relative to neural tissue. We implanted six 32-channel probes, each with two 16-channel shanks (192 of 288 total implanted channels, see Methods) into each of three animals, and recorded continuously (with the exception of moving animal between rooms, see Methods) for 10 or 11 days (animal A, day 53 to 63 post-implant, animal B, day 47 to 57 post-implant, animal C, day 42 to 53 post-implant). Animals performed a spatial navigation task three to four times daily, running ~250 meters during each session. Behavioral sessions were performed in two different rooms. Each 16-channel shank yielded ~1.6 Terabytes of data for that period, and these data were divided into 10 segments of 24-hr length and clustered using MountainSort (Chung et al., 2017). Subsequently, clusters were linked across segments using a simple and conservative mutual nearest-neighbor rule (see Methods and validation in Supplementary Fig. 3a).

This approach allowed us to continuously track a substantial fraction of units across many days, despite the expected waveform variation (Dhawale et al., 2017). An example of a unit that was tracked for the entire period is shown in Figure 4a-d, and on this shank, 24 of 41 clusters identified in the first 24-hour segment could be tracked for more than one week of recording (Fig. 4e). Across the ten shanks (4 from animal A, 2 animal B, 4 animal C), 26% (187 / 707) of clusters could be tracked for 7 days of recording or more (Fig. 4f, Supplemental Fig. 3c), yielding a dataset from these three animals that permits an in-depth analysis of long-timescale changes in single unit activity.

**Figure 4.**
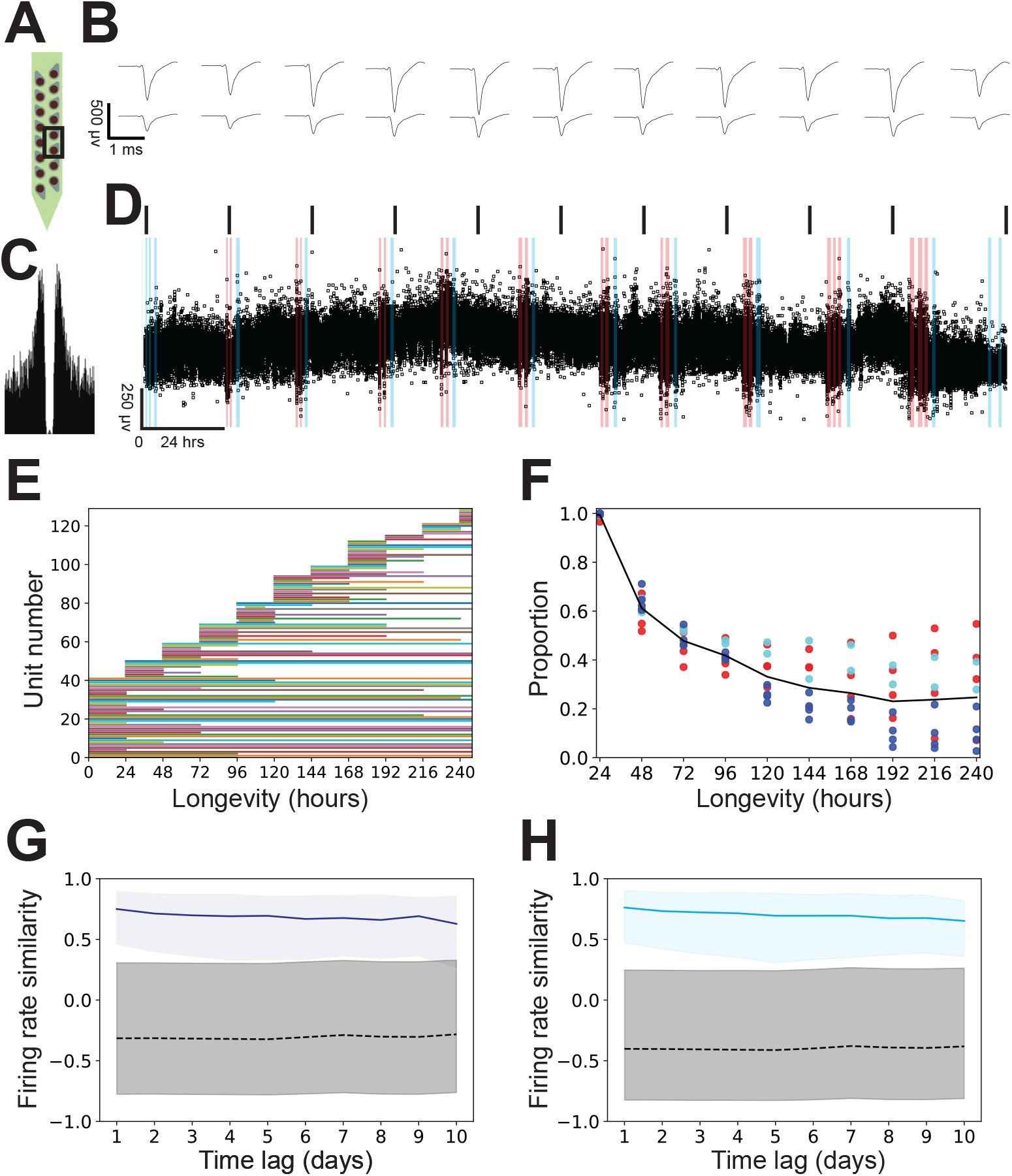
Tracking individual single-units over time. (A-D) Example neuron tracked for 248 hours of continuous recording. (A) Geometric layout of recording channels, with 2 boxed channels on which the unit was clustered. (B) Average waveforms (bandpass filtered 300 – 6000 Hz) for the two channels indicated in (A), calculated for 1-hour time bins every 24 hours, except for the last bin, which corresponds to the last hour of recording (hour 247 to 248). Scale bar corresponds to 400 μv and 1 ms. (C) Autocorrelogram for the unit, calculated over all 248 hours. X-axis corresponds to ± 50 ms in 0.5 ms bins, y-axis normalized to largest bin. (D) Spike amplitude (bandpass filtered 300 – 6000 Hz) over length of continuous recording, for all ~700,000 events in the time period. Each event is shown as a black square, allowing all outliers to be seen. Top, black lines correspond to the 1-hour bins from which average waveforms in (B) are calculated. Shading corresponds to spatial behavioral task performance either in room A (blue), or room B (red, see Methods for more details). Non-shaded times animal was either in rest box or home cage. (E) Period over which each cluster could be tracked for one shank. (See Supplemental Fig. 3 for all other shanks). (F) Proportion of clusters that could be tracked for a given length of time. Black is the total across ten shanks. Each point corresponds to an individual shank from animal A (blue), animal B (cyan), or animal C (red). (G, H) Firing rate similarity for all 3 animals, calculated during behavioral task performance in room one for either (G) low velocity times (< 4 cm / s) or (H) high velocity times (≥ 4 cm / s).

### Firing rate stability in a well-learned task

In the absence of external perturbations, the majority of single-neurons show stable responses when measured intermittently across days (Dhawale et al., 2017; Greenberg and Wilson, 2004; McMahon et al., 2014; Rose et al., 2016). Similar observations have been made from daily recordings in rodent mPFC during spatial behaviors from 60 units across 2 days, and 8 units across 6 days (Powell and Redish, 2014), suggesting that rodent mPFC units show stable firing properties in the context of well-learned behaviors. Our goal was therefore to validate our recording and automated drift tracking methods in comparison to previous findings for rodent mPFC, and to determine whether the observed stability could be confirmed with continuous recordings over longer timescales with a much larger dataset (187 units followed for a week or more).

We focused on a simple measure of unit activity: mean firing rates. Unsurprisingly, units displayed a large range and diversity of firing rates throughout a day (Hromadka et al., 2008; Mizuseki and Buzsaki, 2013; O’Connor et al., 2010). We chose to focus on times where behavior was similar across days, and therefore chose periods when the subjects were performing a well-learned spatial behavior in a familiar environment. The behavioral states were further subdivided into times when the animal was at low (< 4.0 cm / s) and high (≥ 4.0 cm / s) speeds, as these are known to correspond to different neural states (Kay et al., 2016; Yu et al., 2017). For each unit, firing rates were calculated during these times across all ten (n = 2) or eleven (n = 1) days of continuous recording. Importantly, given the large diversity of firing rates between neurons, observing stable single-unit firing rates could only occur if both single-unit physiologic firing rates were stable and the method correctly identified individual cells across time (note here that our spike sorting methodology does not use rate or timing information).

Our findings both validate our unit tracking and confirm that firing rates taken from similar behavioral epochs show remarkable degrees of stability across many days (see Supplemental Fig. 3d for one example animal). We quantified that stability using firing rate similarity (Dhawale et al., 2017) at increasing time lags. We compared the distribution of firing rate similarities of all units that could be tracked for multiple days to the distribution of firing rate similarities for every different cluster pair (i.e. cluster pairs with different cluster ID’s), recorded on the same shank, at the same time lag (see Supplemental Fig. 3e for firing rate similarities for one animal). These analyses confirmed that units’ firing rates were more similar within the same unit than between units across all days of recording for all 3 animals individually (all two-sided Wilcoxon rank sum p < 1.0e-8 low velocity; p<1.0e-5, high velocity), and together (Fig. 4 G, H, all two-sided Wilcoxon rank sum p < 1.0e-24, low velocity; p < 1.0e-27, high velocity).

## Discussion

Electrophysiological recordings provide millisecond resolution information about the activity of neurons, and our system makes it possible to access this information simultaneously across hundreds of neurons within a region, in multiple anatomically distant regions, and to do so for a time period spanning months. We demonstrate large-scale recordings from neurons in three widely separated brain structures, the OFC, the mPFC, and the NAc, yielding a conservative total of 375 well-isolated neurons recorded simultaneously. These recordings allowed us to demonstrate widespread and coordinated activation of all three regions at the time of hippocampal SWR events. Moreover, high quality recordings could be obtained across many months. In addition, our system makes it possible to perform continuous 24/7 recording, and with a simple and conservative linking algorithm we track ~25% of single units across more than a week.

Information processing in the brain is distributed, parallel, and dynamic. In contrast, current experiments often focus on a single region, record from small numbers of neurons, and average over many trials to estimate response functions. While these studies provide key insights into brain function, they cannot capture many of the most central elements of neural computation. Our system provides both high density and modularity to allow for recordings of many units across a set of structures of interest, and longevity and stability to study these units across behavioral states and as they evolve. Our approach is complementary to that of the recently reported Neuropixels probe (Jun et al., 2017), and the combination of features of our system – density, modularity, longevity, and stability, enables experimenters to address fundamental, long-standing questions of brain function.

### Density and Modularity

Neural computations depend on local circuits, distributed circuits within a brain region, and widely distributed circuits located across regions. We therefore developed a technology platform designed to sample many neurons across spatial scales. Our individual polymer arrays consist of multiple shanks, each with 16 closely spaced electrode contacts. This geometry allows us to leverage the single unit isolation achievable when multiple electrodes detect signals from the same neurons (Gray et al., 1995) while making it possible to record across multiple insertion sites in the same brain region. These densities resulted in recordings of up to 45 well isolated single units on a shank and on average one unit per recording electrode when devices were placed in neocortex, permitting study of local circuit dynamics in the neighborhood of a shank and, simultaneously, across shanks in the same brain region.

We demonstrated these capabilities with recordings from 375 units distributed across mPFC, OFC and NAc, selected from 1533 identified clusters. The 4-shank, 64-channel probes used here had a larger contact to edge-of-shank distance than the 2-shank, 32-channel probes, which may have contributed to the higher yield per channel seen with the 2-shank versions (Lee et al., 2017b). These recordings allowed us to identify a subset of SWR-modulated OFC neurons and simultaneous modulations of brain activity during hippocampal SWRs across regions. Recordings from populations of this size make it possible to carry out a number of analyses that are either not possible or very difficult with lower unit counts, including simultaneous comparisons of activity patterns across regions. In this respect only the Neuropixels (Jun et al., 2017) probe offers similar recording densities, and in that case the linear arrangement of sites may limit the density of recordings within a single region.

Here we note that while it is tempting to compare recording yields across devices, these comparisons can only be done fairly if the same spike sorting approach is applied in both cases. We used our recently developed, fully automatic spike sorting package MountainSort (Chung et al., 2017) and applied conservative cluster quality metrics to ensure that we were only including well isolated units. Nonetheless, these per-channel yields are similar to those reported recently for an acute implantation of two Neuropixels probes, where ~370 units per probe were recorded from the 384 active sites. A direct comparison of the yields of chronically implanted Neuropixel probes is difficult because only one chronically implanted probe’s cell yield was reported, which was 127 units 49 days after implantation.

### Longevity and Stability

Experiences drive plasticity in neural circuits, thereby modifying the way they process information. Our system provides the capacity to observe how these changes manifest over the seconds to months during which the network reshapes. We maintained high quality recordings for 160 days across multiple devices and animals, and extended one set of recordings to 283 days with only a slight decline in recording quality. The consistent high-quality recordings for 160 days reported here also exceed those reported for the latest generations of imec devices (Mols et al., 2017), including the immobile, chronically-implanted Neuropixels device, where stable total firing rates and un-curated cluster numbers were reported for recordings spanning 56 days (Jun et al., 2017), although those devices may yield longer recordings than reported.

Finally, we demonstrated stability of recordings that makes it possible to study the same units, 24 hours a day, across at least a week. Using a simple and fully automatic algorithm for matching clustered units across time segments, we could track ~25% of units (187 / 707 from 10 shanks) for seven or more days. Moreover, these units’ firing rates were stable during performance of a well-trained behavior. We note here that our quantification and electrode-drift tracking method provides a conservative estimate of trackable units, and that given the simplicity of our algorithm, it is likely that a more sophisticated approach would allow for even better results. The proportion of units we could track across more than a week is similar to that recently reported for a semi-automatic method applied to data from a 64 channel, 16-tetrode based system, which yielded 19 units per day (Dhawale et al., 2017), less than half of our observed per-channel yield. Paired with the ability to implant more channels in multiple regions, our system will enable the observation of experience- or time-driven changes across distributed neuronal populations.

In summary, our system enables the use of large-scale polymer recording arrays in rats, supporting higher channel counts, cell yields, and longevities. In larger animals, where larger impact forces and brain pulsations are present, flexible polymer will likely match or exceed performance of existing chronic recording technologies. The full 22 mm x 22 mm x 25 mm 1024-ch system should fit into existing primate chambers, making its utilization relatively straightforward.

The implantation platform will benefit from future silicon and polymer process advances, which will potentially enable higher channel counts, lower power consumption, and smaller implant sizes. Beyond pure recording applications, the modular design lends itself to integration with new elements that expand the functionality, such as other recording capabilities (Wassum et al., 2008), circuit manipulations (Tooker et al., 2013; Wu et al., 2015), and computational power for closed-loop applications.

## Author Contributions

J.E.C. and L.M.F. designed the experiments. J.E.C. developed the surgical methodology. J.E.C., H.R.J., J.F., C.G.B., and H.L. collected rodent datasets. J.E.C. analyzed the rodent data. J.E.C., J.F.M., A.H.B., L.M.F., and L.F.G., developed the drift-tracking spike sorting methodologies. J.E.C., V.M.T., S.C., A.C.T., K.Y.L., and L.M.F. designed the polymer array geometries. V.M.T., S.C., A.C.T., and K.Y.L. developed fabrication methodology and fabricated the polymer probes. J.E.C., M.P.K., M.K., D.F.L., and L.M.F. designed the acquisition hardware. J.E.C. and L.M.F. wrote the manuscript with assistance from all authors.

## Acknowledgments

We thank M. Stryker, K. Ganguly, and K. Kay for insightful comments, V. Kharzia and A. Kisel for assistance with histology, and M. Borius for assistance with data acquisition hardware. This work was supported by NINDS grant U01NS090537 to L.M.F and V.M.T. and NIMH grant F30MH109292 to J.E.C. The Flatiron Institute is a division of the Simons Foundation.

## CONTACT FOR REAGENT AND RESOURCE SHARING

Further information and requests for resources should be directed to and will be fulfilled by the Lead Contact, Loren Frank.

## EXPERIMENTAL MODEL AND SUBJECT DETAILS

### Rat

All experiments were conducted in accordance with University of California San Francisco Institutional Animal Care and Use Committee and US National Institutes of Health guidelines. Rat datasets were collected from male Long-Evans rats (RRID: RGD_2308852), 623 months of age, with weights ranging from 500-600 g. All rats were fed standard rat chow (LabDiet 5001) in addition to sweetened evaporated milk for reward during behavioral performance. Rats were ordered from Charles River Laboratories at weights of 300-400 g and 3-4 months of age.

## METHOD DETAILS

### Surgical procedure

Male Long-Evans rats (RRID: RGD_2308852), were implanted with polymer probe(s) at 6-12 months of age. Polymer arrays were targeted to a variety of targets (all coordinates given in millimeters relative to bregma: medial prefrontal cortex (mPFC, including prelimbic and anterior cingulate cortices; ±1.2 ML, +1.5 to +4.5 AP, −2.0 to −4.0 DV, 6-8° from saggital), ventral striatum (VS, primarily nucleus accumbens shell; ±0.7 to +1.9 ML, +0.8 to +1.9 AP, −7.2 DV), orbitofrontal cortex (OFC, primarily lateral orbitofrontal cortex; ±3.5 to 3.7 ML, +2.6 to +3.4 AP, - 4.0 DV), dorsal hippocampus (dHPC, ±2.3 to 2.8 ML, −3.5 to −4.0 AP, −4.0 to −6.0 DV). For some subjects, stimulating electrodes and tetrode microdrives were also implanted at the same time, targeted to the ventral hippocampal commissure (vHC, ±1.0 ML, −1.2 or −2.0 AP) and dHPC.

Anesthesia was induced using ketamine, xylazine, atropine, and isoflurane. Every 4 hours, the animal received additional Ketamine, xylazine, and atropine.

The skull was cleaned, targets were marked, and all drilling was completed. Commercially-pure titanium (CpTi) 0-80 set screws (United Titanium, OH), selected for their well-known ability to osseo-integrate (Le Guehennec et al., 2007), were then placed around the perimeter of the implant. Bone dust was cleared from the skull, and craniectomies and durectomies were completed. The skull was briefly allowed to dry and a custom 3d-printed base piece (RGD837 Stratasys, MN) was then fixed to the skull using 4-META/MMA-TBB (Matsumura and Nakabayashi, 1988) (C&B Metabond). This base piece serves a multitude of functions, including acting a reservoir for saline or silicone gel, an anchoring point for the polymer arrays, and a standardized interface from which the rest of the implant can be affixed and constructed during the implantation.

Polymer probes attached to silicon stiffeners by polyethylene glycol (PEG) were then inserted to the brain (Felix et al., 2013) using custom 3d-printed pieces, avoiding surface vasculature. Polymer probes were then affixed via a piece of polyimide to the 3d-printed base piece before PEG was dissolved using saline, and silicon stiffeners were retracted. Gentle bends were allowed to form below the anchoring points on the polymer arrays, acting as strain relief. Insertion was repeated for all targeted locations.

After all polymer probes were affixed, the saline filling the 3d-printed base piece was then removed and silicone gel (Dow-Corning 3-4680) was used to fill the 3d-printed base piece, providing a means to seal the durectomies and craniectomies, and also provide added support for the polymer arrays. Additional custom 3d-printed pieces were used to construct a protective case around the polymer devices and active electronic components of the implant. Silicone elastomer (Quik-sil, WPI) was then added to the remainder of the exposed polymer, with special attention to the soft polymer – rigid printed circuit board interface, and 3d-printed casing was affixed to the skull using dental acrylic.

### Reagents and data acquisition

#### Polymer arrays

The polymer arrays were fabricated at the Lawrence Livermore National Laboratory nanofabrication facility as described previously (Tooker et al., 2012a, b). Briefly, devices have three trace metal layers and four polyimide layers with a total device thickness of 14 μm.

Devices with an LFP configuration had 20 μm contacts in a single-line with a center-to-center distance of 100 μm, tapered shank width of 61 μm to 80 μm, 21 or 22 contacts per shank, and an edge-of-shank to edge-of-shank distance of 420 μm.

Devices with a 4-shank, 64-channel single-unit configuration are diagrammed in Fig. 1, and had an edge-of-shank to edge-of-shank distance of 250 μm. This design was used in the 1024-channel rat implant, and one module was used in a 352-channel implant (one 4-shank 64-channel module alongside six 2-shank 32-channel arrays, and 24 tetrodes).

Devices with a 2-shank, 32-channel single unit configuration had an identical shank layout to the 4-shank configuration with the notable reduction in edge-of-contact to edge-of-shank distance from 12 μm (4-shank design) to 6 μm (2-shank design). This device design was used for the majority of the data shown, used in the 128-channel implant (data shown in Fig. 3), and all 288-channel implants (six, two-shank, 32-channel polymer arrays and 24 tetrodes).

The device with a 2-shank, 36-channel single-unit configuration (featured in Supplemental Fig. 2) had a similar dual-line, staggered design to the other single-unit configurations with a few notable exceptions. The shank width was 100 μm, edge-of-contact to edge-of-shank distance was 12 μm, and 3 of the 18 contacts were placed closer to the tip of the shank.

#### 16-module, 1024-channel implant

The 16-modules were distributed equally across both hemispheres. Of the 16 modules implanted, 2 were targeted to dHPC and of an LFP configuration. Of the remaining 14 modules, 4 were targeted to OFC, 4 were targeted to VS, and 6 were targeted to mPFC. There were device failures on 4/6 targeted to mPFC, and 2/4 targeted to VS.

#### 160 day periodic recordings

Polymer probes were targeted to mPFC or OFC. In one implant, two two-shank 36-channel arrays were implanted into mPFC and recorded from for 263 days, the termination of the experiment due to animal approaching end of life expectancy. This animal was recorded from using the NSpike data acquisition system (L.M.F. and J. MacArthur, Harvard Instrumentation Design Laboratory) in a 13’’ x 13’’ rest box, and was returned to its home cage. The second implant consisted of four 2-shank 32-channel arrays, all targeted to OFC (128-channel implant). The third animal was implanted with six 2-shank 32-channel polymer arrays targeted to mPFC, alongside two stimulating electrodes targeted to vHC, and 24 tetrodes targeted to dHPC bilaterally, for a total of 288-channels of recording. For the longevity analyses, the second and third animals were also recorded from in a 13’’ x 13’’ rest box, but on some unanalyzed days, recordings were also carried out while the animal ran in a spatial environment.

#### 10-day continuous recording in mPFC

Three animals were implanted with six, two-shank, 32-channel polymer arrays targeted to mPFC, alongside two stimulating electrodes targeted to vHC, and 24 tetrodes targeted to dHPC bilaterally. One of the three animals also had one four-shank, 64-channel polymer array targeted to right OFC. This same animal had a device failure resulting in two functional 32-channel polymer arrays in mPFC and one 64-channel polymer array in OFC. Another animal had a commutator failure on day 4 of recording, causing intermittent data loss, and firing rates from this animal’s day of recording were not used for firing rate analyses. Recordings were carried out while animals were housed in their home cages and in alternating epochs of exposure to a familiar rest box and one of two spatial environments in different rooms. Data was not collected when the animal was being moved between rooms. Animals ran 600 – 1000 meters per day in these spatial environments and provided a challenging experimental setting in which to assess recording stability.

On the first day of continuous recording, animals stayed in one room, room A, where they had been performing the same spatial task for several weeks, and performed three behavioral sessions, each lasting 30 – 40 minutes. On the second day of recording, animals performed two 30 – 40 minute behavioral sessions in room B, their first time being exposed to that room, and then one in room A. On days three through eleven, animals performed two or three sessions of behavior in room B followed by one in room A. Recording was stopped half an hour after the animal finished the session of behavior in room A on day eleven (animals A and B), or day twelve (animal C). In animal C, a twelfth day of recording was carried out with all behavioral sessions occurring in room A. Animals had red/green tracking LED arrays attached to the implant, allowing their position to be extracted from video recorded by a camera mounted to the ceiling.

### Data processing and analysis

Data analysis was performed using custom software written in Python 3.6.3 (Anaconda linux-64 v7.2.0 distribution, Anaconda Inc.) and Matlab 2015b (Mathworks).

#### Spike sorting

Clustering was done using MountainSort, using settings and thresholds as reported previously (Chung et al., 2017). Adjacency radius was set to 100 μm when sorting the 20 μm contact, 20 μm edge-to-edge dual-line designs, resulting in clustering neighborhoods of 5 to 9 electrodes. The event detection threshold was set to 3 SD. Putative single-units were identified using previously set thresholds (isolation > 0.96, noise overlap < 0.03) and an automatic merging procedure, reported previously (Chung et al., 2017), was used to identify pairs of clusters that corresponded to the higher and lower amplitude components of single units that fired in bursts with decrementing spike amplitudes.

For the 240-hr continuous recording datasets, filtering and spatial whitening was applied to the entire 240-hr recording, and then data was clustered in 24-hour segments. Automated curation and bursting-related merging was first completed independently for each segment. As a result, all clusters in all segments satisfied our criteria for well isolated units. Linking clusters between segments was done using a mutual nearest neighbor rule. For every cluster in the first segment, a 1.66 ms spatially-whitened waveform template was calculated from the last 100 events, using every channel on the shank. Similarly, for every cluster in the second segment, a waveform template was calculated from the first 100 events. Next, the L^2^ distance was calculated between every segment 1 and segment 2 pair of templates. If cluster A from segment 1 and cluster A’ from segment 2 were mutual nearest neighbors, then the segments were linked.

This approach is conservative as a result of three main features. First, it used only well isolated clusters from each segment, and only matched these well isolated clusters. Second, because the 24-hour segments were not aligned to specific events in the animals’ experience, the segments partitioned the spiking activity at points where large, sudden changes in spike amplitudes were very unlikely. Third, the distance calculation was based on whitened spike waveforms from the entire 16 electrode array, yielding unique templates for each unit. The mutual nearest neighbor calculation ensured that these templates matched across the segment boundaries, and we found that this linking algorithm yielded plots of spike amplitude over time that were continuous across the period where the unit could be tracked.

#### SWR detection and modulation

SWRs were detected as previously described (Cheng and Frank, 2008). Briefly, LFPs from a contact near CA1 was filtered into the ripple band (150 – 250 Hz) and the envelope of band-passed LFPs was determined by Hilbert transform. SWR were initially detected when the envelope exceeded a threshold (mean + 3 SD) on the contact. SWR events were defined as times around the initially detected events during which the envelope exceeded the mean. For SWR-trigged firing rates, only SWRs separated by at least 500 ms were included.

SWR modulation analysis was carried out as described previously (Jadhav et al., 2016). Briefly, spikes were aligned to SWR onset resulting in SWR-aligned rasters. Cells with less than 50 spikes in the SWR-aligned rasters were excluded from these analyses. To determine the significance of SWR modulation, we created 1,000 shuffled rasters by circularly shifting spikes with a random jitter around each SWR, and defined a baseline response as the mean of all shuffled responses. We then compared the response in a 0-200 ms window after SWR onset (SWR response) to the baseline. We considered a cell as SWR-modulated when the mean squared difference of its shuffled response from the baseline (i.e., p < 0.05). SWR-modulated neurons were further categorized as SWR-excited or SWR-inhibited by comparing the rate in a 0-200 ms window after SWR onset, with the rate of the mean shuffled response in the same 0-200 ms window.

#### Generalized linear models during SWRs

Construction of generalized linear models (GLMs) was done as reported previously (Rothschild et al., 2017). Briefly, the GLMs were constructed with a log link function to predict spike counts of single units during SWRs in PFC, NAC, or OFC from ensemble spiking patterns in another region. The region’s SWR ensemble pattern was the vector of binned spiking responses across units recorded in that region during the 0-200 ms window after SWR onset.

The ensemble patterns were used to predict single cell SWR responses. A single prediction model was generated using predictor data of the ensemble patterns across SWRs, and predicted data of the single-cell SWR responses across SWRs. Only cells that were active (> 0 spikes) in more than 10 SWRs were predicted. For each predictor ensemble and predicted cell, we performed five-fold cross validation. We randomly partitioned the SWRs into five equally sized sets, with the constraint that the number of nonzero values in the predicted vector must be approximately balanced across sets. For each fold, four of five folds was used to train the GLM, and the remaining fold to test. For the test phase, the model derived from the training phase was applied to the predictor ensemble data in the test set, yielding predictions for the predicted cell firing across SWRs.

Prediction error was defined as the mean absolute difference between the predicted spike counts and the real spike counts. For that same fold, we defined a baseline prediction error by performing 100 random shuffles of the predicted firing rates across SWRs in the test fold and taking the mean of the shuffled prediction errors. The real and shuffled prediction errors were then averaged across the five folds. Prediction gain for one predictor-ensemble-predicted-cell combination in one time window was defined as the shuffled prediction error divided by the real prediction error.

For comparison, we repeated the exact same procedure described above on 100 random shuffles of the entire original dataset, where shuffling entailed random matching of activity patterns in the predictor and predicted data (*e.g.*, taking predictor data from one SWR and using it to predict firing rate for another SWR). To assess prediction significance for a pair of regions, we compared the distribution of real prediction gains to the shuffled prediction gains across all ensemble/cell combinations using a two-tailed nonparametric Wilcoxon rank sum test.

#### Cluster linkage analysis

Quantification of the relative distances of successfully linked cluster pairs to the other possible linked clusters (Fig. 4F) was done as follows: if there was a successful link made between cluster A from segment 1 and cluster A’ from segment 2 (A to A’), then the L^2^ distances between cluster waveform templates (A and B’), (A and C’),…(B and A’), (C and A’), etc., were normalized to the L^2^ distance of (A to A’). These distances, for all successfully linked pairs across all electrode arrays, contributed to the histogram in Fig. 4F.

To quantify the distances of successfully linked cluster pairs and their distance to other possible linked clusters relative to the variability of the events within the successfully linked cluster, we normalized the same set of distances as above using the mean spike distance to its template. Specifically, if there was a successful link made between cluster A from segment 1 and cluster A’ from segment 2 (A to A’), the mean of the L^2^ distances between the 100 events and the template of A (calculated from the same 100 events) was used as the normalization factor for the L^2^ distance from (A to A’), and all other unlinked pairs, (A and B’), (A and C’),…(B and A’), (C and A’), etc. This mean of the L^2^ distances is referred to in the text as “event distance.”

In Fig. 4G, the normalized distances of successful linkages, (A to A’), contributed to the histogram in red, while the normalized distances of all other unlinked pairs, (A and B’), (A and C’),…(B and A’), (C and A’), etc., contributed to the histogram in black.

#### Firing rate similarity during behavioral performance

Firing rates were calculated for when the animal was performing the spatial behavior in room A. This constituted ~90 minutes of time on day one (and day twelve in animal C), or ~30 minutes of time on days two through eleven. Roughly half of the time during behavioral performance was spent either at low (< 4.0 cm / s) or high (≥ 4.0 cm / s) velocities.

Firing rate similarity was calculated using the same formula as in (Dhawale et al., 2017), where the similarity of two different firing rates, *FR_i_* and *FR_j_* was measured by the following formula:

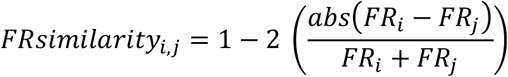

A firing rate similarity score of 1 occurs when *FR_i_* and *FR_j_* are identical, and a firing rate similarity score of −1 occurs when one firing rate is 0 (maximally dissimilar). As in (Dhawale et al., 2017), when comparing firing rates for the same unit across time, firing rate similarity was calculated for time lags ranging from 1 to 10 days (animals A and B), or 11 days (animal C, Supplemental Fig. 3). In other words, if a cell could be tracked for all 12 days of behavioral performance in room A, its 1-day time lag firing rate similarity was calculated 11 times (days 1-2, 2-3,…10-11, 11-12), or its 10-day time lag was calculated twice (days 1-11, 2-12).

The distribution of within-unit time lagged similarities was compared to the distribution of all between-unit time lagged similarities, matched for both shank and time lag. This differs from the comparison done in (Dhawale et al., 2017), where time-lagged similarities were compared to the within-day across-unit distribution of firing rate similarities.

### Data availability

The data that support the findings of this study will be made available upon reasonable request.

### Code availability

Electrode-drift spike sorting code is available at https://github.com/magland/msdrift. This code is designed to be a package added to the core MountainSort software, available at https://github.com/flatironinstitute/mountainsort. The analysis code used in this study will be made available upon reasonable request.

## Supplementary Figure Legends

**Supplemental Figure 1.**
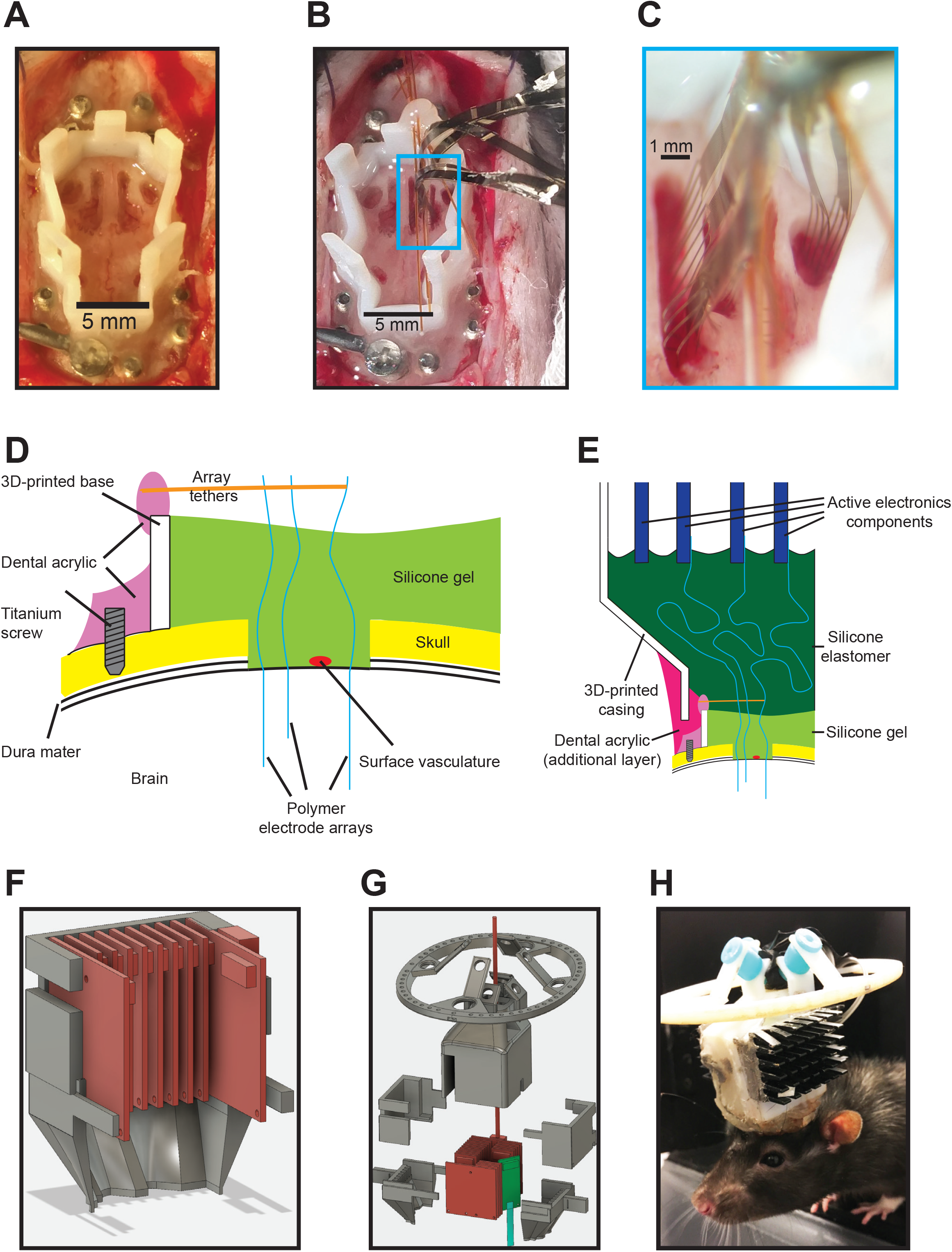
Surgical approach and implant construction. (A, B) Top-down views of a rat skull with 3-D printed implant base attached (A) before polymer array insertion, and (B) after insertion of 7 polymer probes. (C) Magnified view of polymer probes entering into brain. (D) Cross-sectional schematic of implant after arrays have been inserted and silicone gel has been added to the 3-D printed base, and (E) of the assembled implant, with silicone elastomer fill to protect soft passive electrical components and moisture-sensitive active electrical components, and to provide strain relief for their soft-hard interface. (F) 3-D model of active electronics (red) and casing (grey), which provide structural support and protection for the passive electrical connection from the implanted contacts to the active electronic components. (G) 3-D model of full implant with polymer probe (cyan), single 64-ch board module (green), active electronics and micro-HDMI cable (red). (H) Rat implanted with full system, including heat sinks (black) and silicone grommets for impact resistance (cyan).

**Supplemental Figure 2 (corresponding to Fig. 3).**
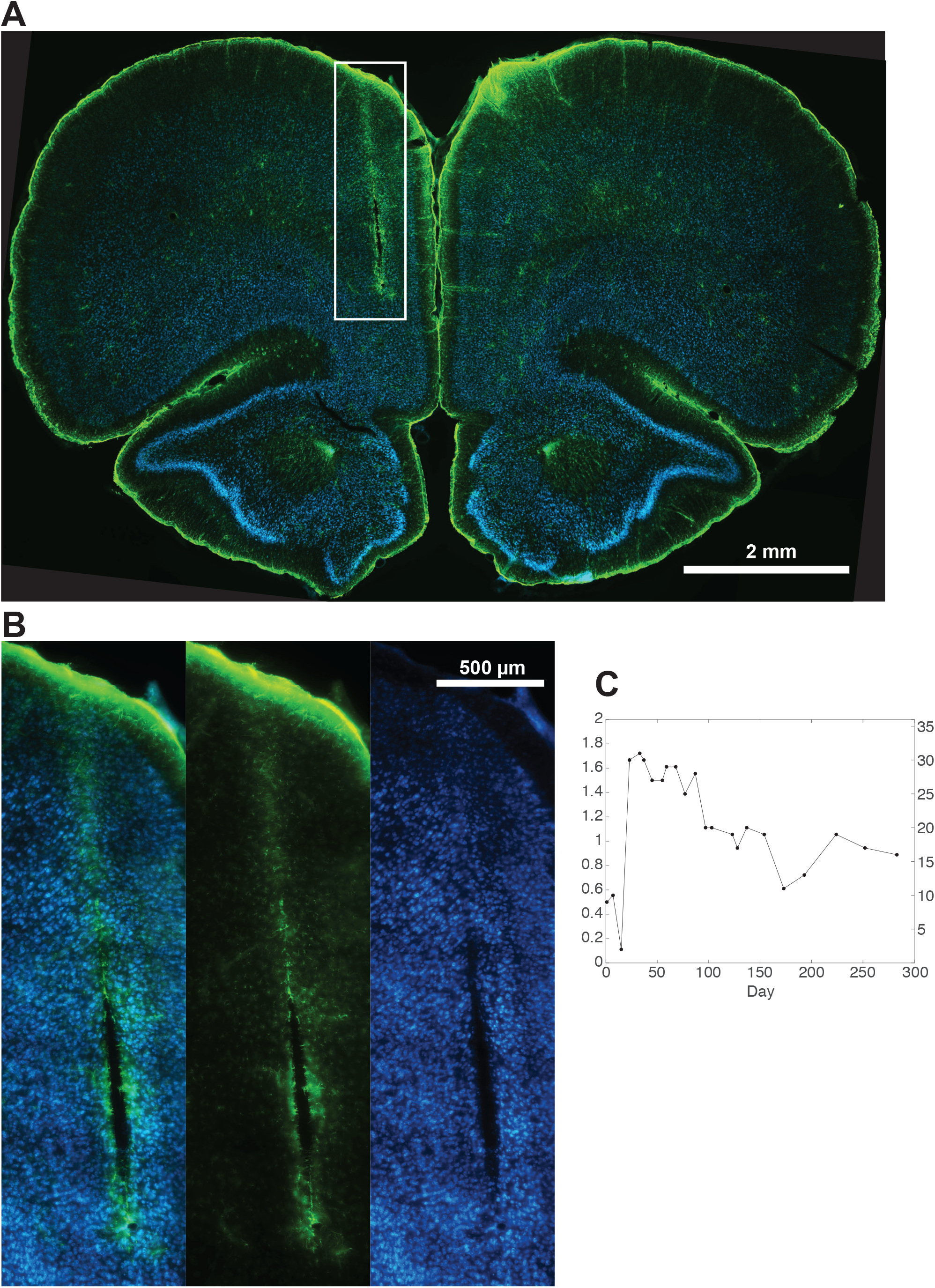
Histology 160 days after implantation. Histology shown corresponds to shank with green arrowhead in Fig 3A. (A) Merged image with glial fibrillary acidic protein (GFAP) stain in green, and NeuroTrace (ThermoFisher Scientific) in blue (B) As in (A), but for highlighted region. Left, merge, middle, GFAP, right, NeuroTrace. (C) Cell yields per channel (left y-axis) or per 18-ch shank (right y-axis) for a probe implanted for 283 days. Experiment was terminated due to animal approaching end of expected lifespan.

**Supplemental Figure 3 (corresponding to Fig. 4).**
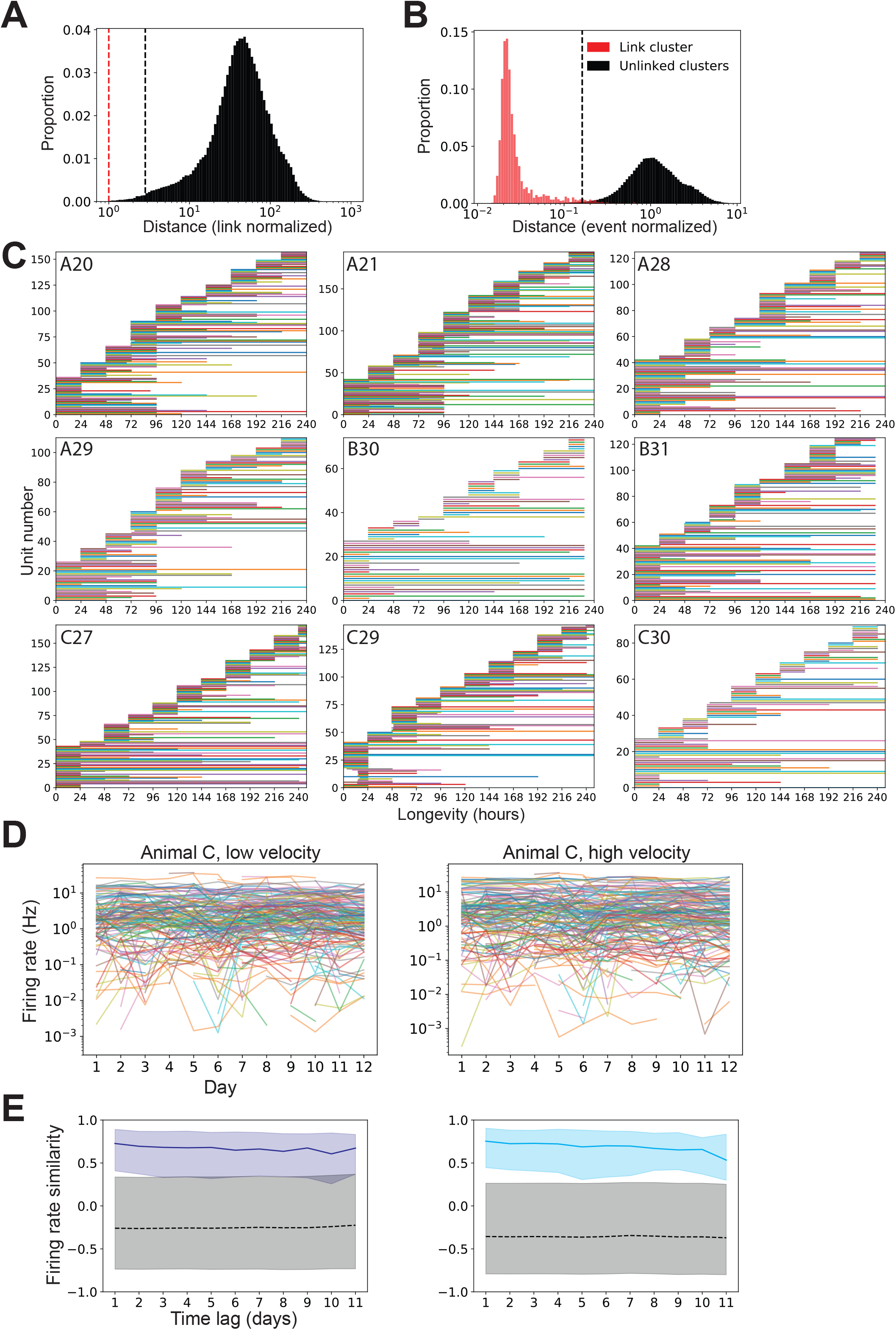
Validation of cluster linkage and stability of single units. (A) If the clusters from each segment were drifting to a greater degree than the separation between clusters, the mutual nearest neighbor cluster pairs could occur in a crowded feature space, with unlinked clusters lying close to the linked cluster. This would generate an environment where erroneous linkages could be made, causing an overestimation of how stable clusters were. To validate that the linkages between 24-hour segments were occurring in cases where the mutual nearest neighbors were unambiguous the distances between linked cluster template to all other possible linking cluster templates (n = 254,034), normalized by the distance between the two linked cluster templates (n = 2,962) were calculated. Shown is a histogram of these distances, where the vertical red line marks unity, the distance of all linked cluster templates. Over 99% of all other possible linking templates lie to the right of the vertical black line (2.8 times the distance to the linked template). (B) When a cluster is stable, the variability of the events should be larger than the change in the template over time. To confirm that the clusters being linked fell within the variability of events around the cluster, we normalized the cluster pair distances by the mean distance of the last 100 events in a cluster from its template (“event distance”, see Methods for more details). Shown is a histogram of distances as in (A), with distances between linked cluster templates (red, n = 2,962), and linked cluster to unlinked cluster templates (black, n = 254,034), but instead normalized by the average distance of the last 100 events from their template. Over 99% of all other possible linking templates lie to the right of the vertical dotted black line (0.16), while 97.5% of linkage distances lie to the left of the vertical dotted black line. The distance between all linked cluster templates was less than their respective within cluster event distances (all < 1). (C) Period over which each cluster could be tracked, separated by inset shank id. (D) Firing rates of all clusters from animal C while performing the spatial behavioral task in room A during either low velocity times (<4 cm / s, left) or high velocity times (≥ 4 cm / s, right). (E) Firing similarities at different time lags, calculated from firings rates shown in (D), from animal C while performing the spatial behavioral task in room A during either low velocity times (<4 cm / s, left) or high velocity times (≥ 4 cm / s, right).

